# Specific phenotypic, genomic, and fitness evolutionary trajectories toward antibiotic resistance induced by pesticide co-stressors in *Escherichia coli*

**DOI:** 10.1101/2021.07.13.452255

**Authors:** Yue Xing, Xiaoxi Kang, Siwei Zhang, Yujie Men

## Abstract

To explore how co-occurring non-antibiotic environmental stressors affect evolutionary trajectories toward antibiotic resistance, we exposed susceptible *Escherichia coli* K-12 populations to environmentally relevant levels of pesticides and streptomycin for 500 generations. The coexposure substantially changed the phenotypic, genotypic, and fitness evolution trajectories, resulting in much stronger streptomycin resistance (>15-fold increase) of the populations. Antibiotic target modification mutations in *rpsL* and *rsmG*, which emerged and dominated at late stages of evolution, conferred the strong resistance even with less than 1% abundance, while the off-target mutations in *nuoG, nuoL, glnE*, and *yaiW* dominated at early stages only led to mild resistance (2.5 – 6-fold increase). Moreover, the strongly resistant mutants exhibited lower fitness costs even without the selective pressure and had lower minimal selection concentrations than the mildly resistant ones. Removal of the selective pressure did not reverse the strong resistance of coexposed populations at a later evolutionary stage. The findings suggest higher risks of the selection and propagation of strong antibiotic resistance in environments potentially impacted by antibiotics and pesticides.

## Introduction

Antibiotic resistance poses a major threat to public health worldwide. It has been estimated that 700 000 people died every year due to antibiotic-resistant bacterial infections during 2014 and 2016, globally^1^. This number is even predicted to reach as many as 10 million by the year of 2050 if no actions are taken^1^. In addition to the increase of antibiotic-resistant pathogens in clinical settings, antibiotic-resistant bacteria and resistance genes are frequently detected in natural and engineered environments, such as surface water, soil, wastewater, and sludge^2-5^. Resistant bacteria developed in those environments may re-enter the water cycle and food chain, potentially imposing health risks. Thus, it is urgent to control the emergence and spread of antibiotic resistance in those environmental hotspots.

The resistance of the microorganisms could either be obtained by *de novo* mutations through evolution or mediated by horizontal transfer of resistance genes. The evolution of antibiotic resistance is driven by selective pressures. Antibiotics are the long-standing focus to study selective pressures. Antibiotics, both at lethal levels and below minimal inhibitory concentrations (sub-MIC) are able to facilitate resistance evolution, including the selection of *de novo* resistant mutants and the selection of preexisting resistant mutants over a long-term selection^6-11^. The two selection levels of antibiotics can induce specific genetic mutations, which render different resistance mechanisms to microbial populations^10-12^. With lethal levels of antibiotics, cells either die or survive, depending on the spontaneous acquisition of resistance-specific mutations, which in general confer high levels of resistance. In contrast, antibiotics at sub-MIC levels do not kill cells and tend to favor more diverse mutations, the combination of which may also cause strong phenotypic resistance^10^.

The resistance developed at sub-MIC levels raises more environmental concerns because in many environments, such as surface water, wastewater, biosolids, agricultural soils, and surface runoffs^13-19^, antibiotics occur at sub-MIC levels. However, in most of those environments, antibiotic residues do not exist alone. Other organic contaminants, such as non-antibiotic drugs, personal care products, and pesticides, often coexist with antibiotics^19-21^. Nevertheless, co-selection by two or more stressors remains poorly understood. Our recent study reveals a synergistic effect of pesticides on the development of resistant mutants under the selection of sub-MIC ampicillin^18^. As a result, the evolution of antibiotic resistance could have been underestimated in environments with the presence of both antibiotics and non-antibiotic chemicals. This study also raised more fundamental questions on how co-stressors would shape a bacterial population during a long-term evolution and how mutational dynamics would affect the phenotypes in terms of antibiotic resistance and growth fitness.

The process of antibiotic-resistance evolution could be divided into the emergence of resistant mutants, the proliferation and maintenance of resistant subgroups in a bacterial population. Previous studies that identified stressors selecting antibiotic resistance mainly focus on resistant mutants and mutations developed at the endpoint of the evolution. These studies facilitated the understanding of the interplay between stressors and the most beneficial mutations. However, this type of study fails to demonstrate the sustainability of the resistant lineages in a population, the dynamics of genetic adaptation, and the associations of resistance evolution with genome and fitness evolution. All of these factors are of great importance to understand the spread and persistence of resistant bacteria in their receiving environments, as well as to predict and intervene in the evolution of antibiotic resistance.

In this study, we aimed to fill the knowledge gap and conducted laboratory evolution experiments by exposing susceptible *Escherichia coli* strain K-12 to sub-MIC streptomycin (Strep) and non-antibiotic organic pollutants for 500 generations. We chose Strep as the exposed antibiotic because Strep is one of the human antibiotics that have been widely used in plant agriculture to combat bacterial diseases like the citrus greening disease^22^, highly likely co-occurring with pesticides and non-antibiotic drugs. Instead of endpoint evaluation of resistance development, we focused on the trajectories of Strep resistance, genomic, and fitness evolution. We investigated the associations of evolutionary trajectories of Strep resistance with the trajectories of genetic mutations and growth fitness in the coexposed populations. Stimulated development of strong Strep resistance by the non-antibiotic co-stressors, pesticides but not pharmaceuticals, was demonstrated. Novel mutations leading to Strep resistance were identified, which expands our fundamental knowledge of antibiotic resistance mechanisms developed under environmentally relevant exposure conditions. Moreover, relative fitnesses of the identified resistant mutants in the coexposed populations were determined, which could be used to predict the proliferation and spread of certain resistant mutants once they are transported into a different environment.

## Materials and Methods

### Bacterial strains, growth, and evolutionary experiments

The bacterial strain used in this study was Gram-negative *Escherichia coli* strain K-12 (ATCC. 10798), which has been widely used as the susceptible and negative control in many relevant studies^23-27^. The growth medium for all evolutionary experiments was Luria-Bertani (LB) broth. First, the stock *E. coli* cells were revived and then streaked on an LB agar plate. After 20-hour incubation, one single colony was picked and inoculated into LB broth, which was regarded as the ancestor (G0), and used as the inoculum for the evolutionary experiments.

The selective pressures included a combination of Strep and pesticides, Strep and pharmaceuticals, pesticides only, pharmaceuticals only, and Strep only. The concentrations of Strep included 1/5 MIC_0_ (the MIC of G0, 8 mg/L), which represented a low-level antibiotic selection. The pesticides and pharmaceuticals selected in this study (Table S1) were those frequently detected in various environments. Their concentrations were based on their detected concentrations in the environment (for pesticides: 0.1 – 4.8 µg/L each and ∼20 µg/L in total). Three pesticide/pharmaceutical exposure levels were included, i.e., 1×, 10×, and 100× environmental concentrations [denoted (1/5Strep,1P), (1/5Strep,10P), and (1/5Strep,100P) for pesticides]. The control groups without chemical exposure were also set up using eight independent populations. Moreover, to take into account the potential dose-effect caused by the additional amount of pesticide co-stressor added to the culture besides the primary stressor (1/5MIC_0_ Strep), we included a condition of (1/2Strep,0), where the concentration of Strep (4 mg/L) was comparable to the total concentration (3.6 mg/L) of (1/5Strep,100P). Cell growth under different exposure conditions was measured by optical density at 600 nm (OD_600_) (See Supplementary Methods in the Supplementary Information for details)

Evolutionary experiments were performed as described in our previous study^28^. We serially passaged eight replicate populations of each condition in LB media containing certain concentrations of pesticides/pharmaceuticals and Strep (Fig. 1). During each transfer, the cell culture was first diluted ten times, and then a volume of 4 µL diluted inoculum was inoculated into each well (500× dilution), making the total volume of 200 µL. The plate for each transfer was incubated at 30 °C in a 150-rpm shaker in the dark every 24 hours. The exposure was conducted for about 500 generations (∼ 55 passages). The cultures after every 100 generations were preserved by adding 100 µL of 50% glycerol and stored at −80 °C.

**Fig. 1.**
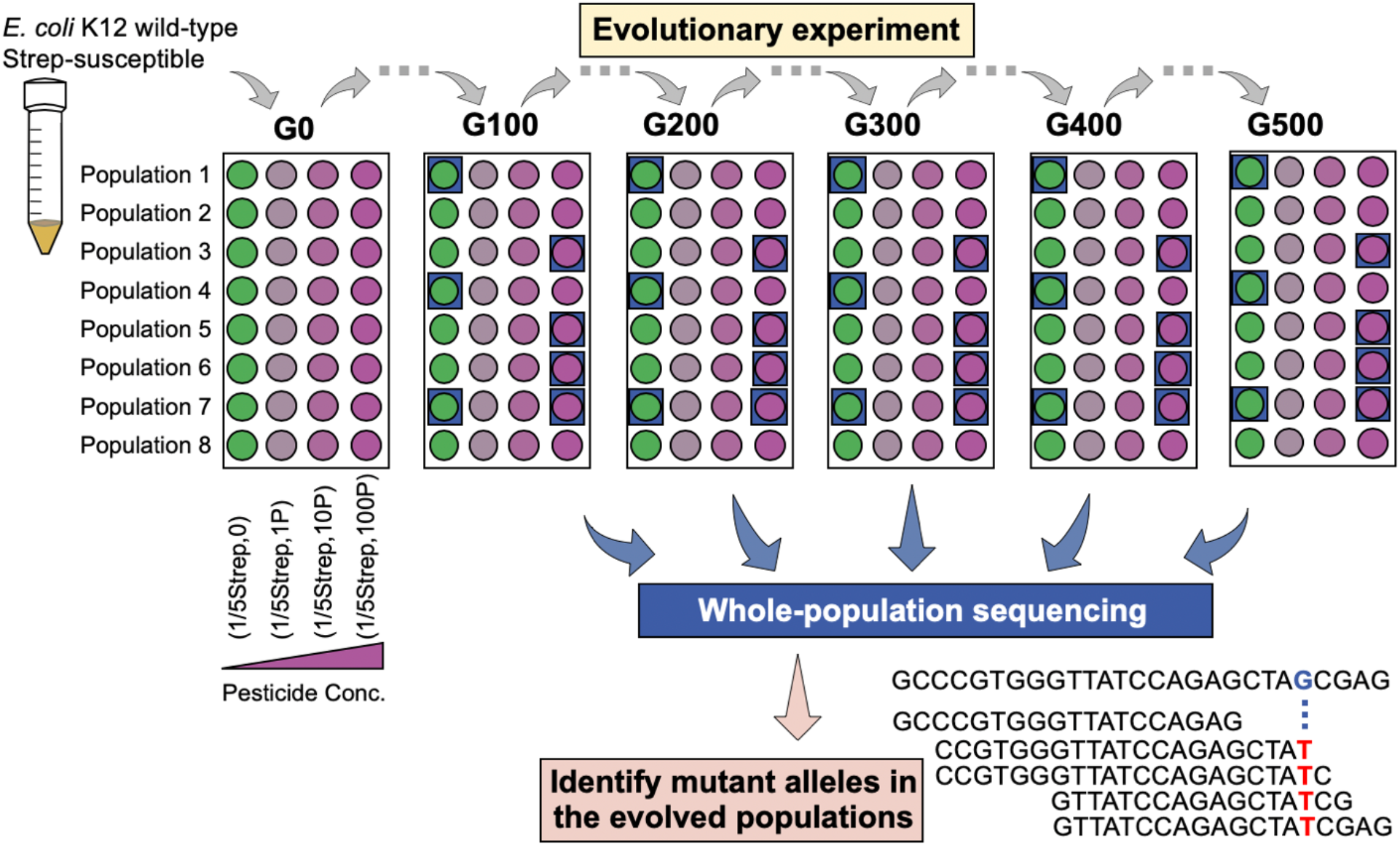
Illustration of the experimental design. A total of 8 parallel populations of *E. coli* K-12 under each exposure condition were serially passaged every 24 h (dilution factor = 1:500, ∼9 generations) into fresh LB medium containing Strep and/or pesticides at the same exposure levels for 500 generations. The exposure conditions include 1/5 MIC_0_ of Strep only, denoted “(1/5Strep,0)”; 1/5 MIC_0_ of Strep and the pesticide mixture: (1/5Strep,1P), (1/5Strep,10P), and (1/5Strep,100P); pesticides only: (0,1P), (0,10P), and (0,100P); and the no exposure control: (0, 0). “P” represents environmental concentrations of the pesticides as listed in Table S1, the total of which was about 20 µg/L. Populations after every 100 generations (highlighted in blue boxes) were subject to the whole-population sequencing.

### MIC test of evolved populations

Every 100 generations, the evolved populations were subject to MIC tests, which determine phenotypic resistance levels of the populations. The cell culture was diluted with 0.9% NaCl solution to an OD_600_ of 0.1, which was regarded as the standard solution. Then 0.5 μL of the standard solution was added into fresh LB medium containing Strep with a series of concentrations. A growth control without the antibiotic and a negative control without bacterial inoculum were set up in the meanwhile. Cell cultures were incubated at 37 °C for 20 hours, and then the OD_600_ was measured. The MIC was determined as the concentration that inhibited 90% of growth based on the OD_600_ measurement^8,29^.

### Determination of population MICs with different fractions of strongly resistant mutants

To determine the correlation between the population MICs and fractions of strongly resistant mutants, we constructed mock populations by growing the mutant and wild-type cells at various ratios. The tested fractions of the resistant mutant in the mock populations included 0, 10^−5^, 10^−4^, 10^−3^, 10^−2^, 10%, 20%, 30%, 40%, 50%, 60%, 70%, 80%, 90%, 99%, 1. These populations were then subject to the MIC test, as described above.

### DNA extraction, whole-population sequencing, and SNP calling

To study the differences in mutational dynamics caused by the addition of pesticides, we selected three lineages from (1/5Strep,0) and four lineages from (1/5Strep,100P). For each lineage, the archived population at every 100 generations was cultivated overnight in LB medium, and cell pellets were collected by centrifugation. Genomic DNA (gDNA) was extracted using the DNeasy Blood and Tissue Kit (Qiagen) according to the manufacturer’s instructions. The gDNA concentration and quality were determined on a Qubit 4 Fluorometer (Thermo Fisher Scientific, Wilmington, DE). The gDNA was then subjected to Illumina NovaSeq 150-bp paired-end sequencing carried out by Roy J. Carver Biotechnology Center at the University of Illinois. The average coverage was 13.3M paired-reads per sample. A dynamic sequence trimming was done by SolexaQA software^30^ with a minimum quality score of 30 and a minimum sequence length of 50 bp. All samples were aligned against the *E. coli* K-12 MG1655 genome available at NCBI GenBank (NC_000913.3) using the Bowtie 2 toolkit^31^. SAMtools was used to format and reformat the intermediate alignment files^32^. SNPs and INDELs were identified and annotated with software BCFtools^33^ and EnpEff^34^. The valid mutant alleles in the sequenced populations were those with (i) amino-acid-sequence change, (ii) not found in the ancestor G0, (iii) >8-read coverage, and (iv) >10% mutant allele frequency at the mutation positions.

### Genotype confirmation

The genotypes of resistant mutants were confirmed by SNP genotyping assays (See Supplementary Methods for details) and whole-genome sequencing analysis. We first isolated resistant mutants in the evolved populations onto selective LB agar plates (1× MIC_0_ Strep), including (1/5Strep,100P)-3 at G300, (1/5Strep,100P)-5 at G400, (1/5Strep,100P)-6 at G300, (1/5Strep,100P)-3 at G500, and (1/2Strep,0)-1 at G200. Three resistant mutant colonies were picked up from each population and subjected to gDNA extraction. The gDNA was tested for the presence/absence of the six mutant alleles. As no successful SNP assays could be designed for the *yaiW* and *rsmG* mutations that were also likely responsible for the resistance phenotype, the genotypes of the resistant mutants isolated from (1/5Strep,100P) populations and those from (1/2Strep,0) populations were further confirmed by whole-genome sequencing following the same procedure as the whole-population sequence described above.

### Competition tests

The competition tests were first carried out in pairs of wild type vs. the mildly resistant mutants, wild type vs. the strongly resistant mutant, and mildly resistant mutant vs. the strongly resistant mutant. Mock populations were constructed by inoculating two types of cells at a ratio of 1:99 and 99:1, separately. Six parallel populations were inoculated for 50 generations under the selection of (1/5Strep,100P) or no selective pressure. The fractions of the two types of cells in the population at the end of cultivation were then determined by comparing the allelic discrimination plots from SNP genotyping assays to those of standard mixtures (Fig. S1). The minimal selective concentration (MSC) of resistant mutants were determined (See the Supplementary Methods).

We then compared the relative fitness of resistant mutants from (1/5Strep,100P) with those from Strep only selections, including (1/5Strep,0)-1 at G500, (1/5Strep,100P)-4 at G500, (1/5Strep,100P)-7 at G500, (1/2Strep,0) at G200. The ratio of competition pairs is approximately from 1:1, and the fraction of (1/5Strep,100P) mutants after 18 generations was determined by SNP genotyping assays. The relative fitness was calculated according to the selection coefficient of cell A/cell B: ln[R(t)/R(0)]/t, as previous described^35^, where R is the ratio of cell A to cell B, t is growth generation. The selection coefficient >0 stands for higher fitness of cell A to cell B. We also used this system to evaluate the individual contribution of pesticides and 1/5 MIC_0_ to the selection of preexisting mutants.

## Results

### Trajectories of phenotypic resistance to Strep in *E. coli* populations with and without pesticide co-stressors

The exposures to Strep at 1/5 MIC_0_ (i.e., 1/5Strep) and/or the pesticides at the highest level (i.e., 100P) (Fig. 1) did not significantly inhibit cell growth, with only 1.7 – 2.9% decrease in the maximum growth rate (Table S2). However, the coexposure to pesticides under 1/5Strep selective pressure substantially altered the evolutionary trajectories of the *E. coli* populations toward high-level resistance. The resistance trajectories of (1/5Strep,0) over 500 generations revealed similar trends among the eight parallel populations, which only acquired low-level Strep resistance with 2.5 – 4× increase in MIC (Fig. 2A). In contrast, when exposed to even 1P pesticides together with 1/5Strep, one out of the eight populations acquired much stronger resistance (≥15× increase in MIC) (Fig. 2B). The other seven (1/5Strep,1P) populations showed similar evolutionary trajectories of Strep resistance as those of the (1/5Strep,0) populations. We observed a dose-effect of pesticide co-stressors on the stimulated evolution toward Strep resistance. More coexposed populations became strongly Strep-resistant as the pesticide level increased, i.e., 2/8 and 4/8 populations exhibited strong resistance (i.e., ≥15× increase in MIC) for the (1/5Strep,10P) and (1/5Strep,100P) exposures, respectively (Fig. 2C&D). The strong Strep resistance can emerge from 200 generations after the coexposure (1/5Strep,1P), where the pesticide concentration (20 µg/L) was much lower than Strep (1.6 mg/L) (Fig. 2B).

**Fig. 2.**
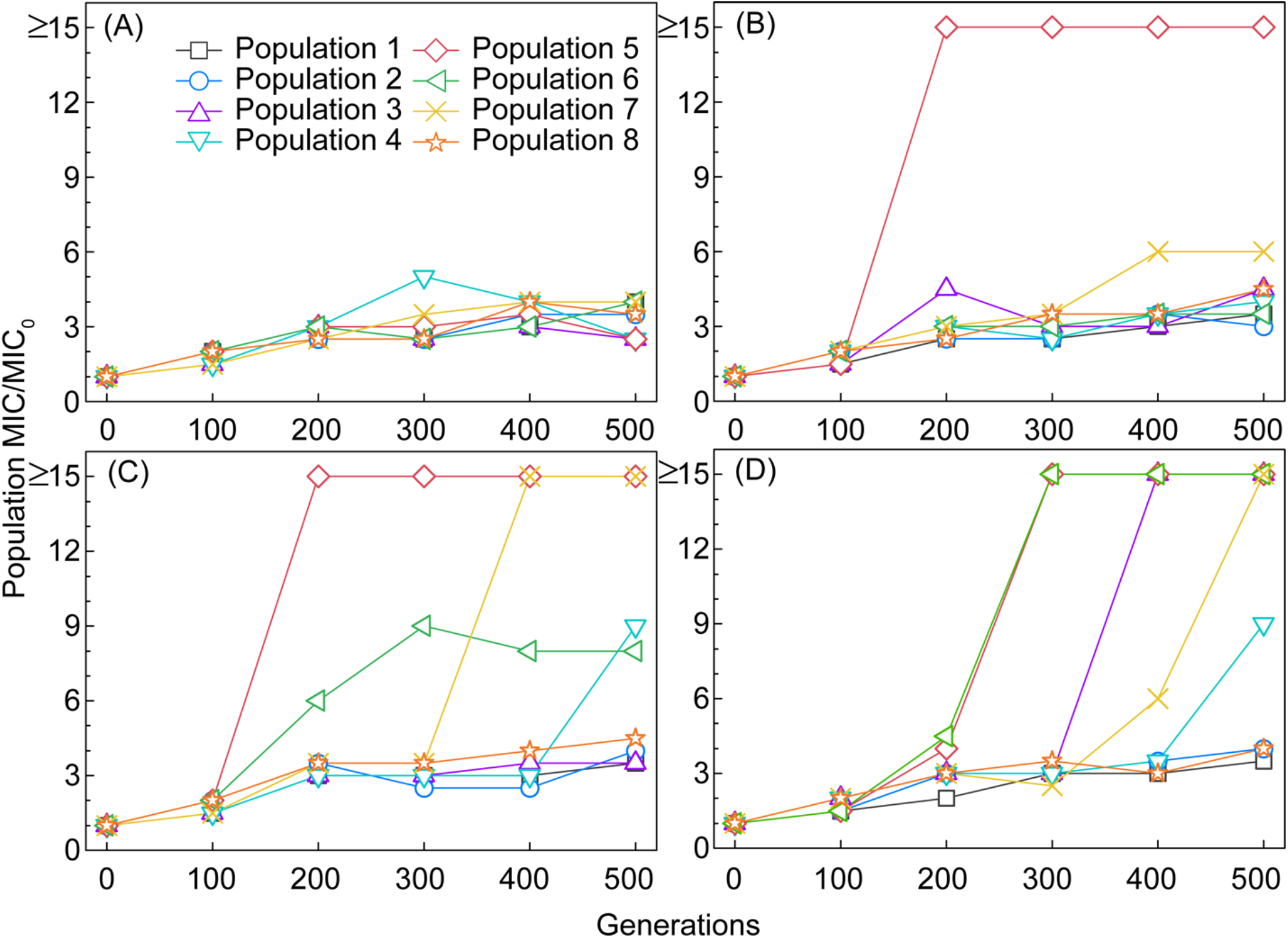
Population MICs to Strep over 500 generations under different exposure conditions (A: (1/5Strep,0); B: (1/5Strep,1P); C: (1/5Strep,10P); D: (1/5Strep,100P)).

Pesticide-only exposures caused ∼ 1.5× increase in MIC in a portion of the eight populations (Table S3). The combined effect of pesticides and Strep in selecting for resistance is much greater than the sum of individual effects of pesticides or Strep, which suggests a synergistic effect. Moreover, the coexposure to 1/5Strep and selected pharmaceuticals (Table S1) did not exhibit a significant difference in MIC increase from that acquired under the exposure to 1/5Strep only (Table S4). Comparing the results of the addition of pesticides and pharmaceuticals, we demonstrated the specificity of pesticides as co-stressors in significantly promoting the evolutionary trajectories toward high-level Strep resistance.

### Distinctive mutational dynamics along the resistance evolutionary path in *E. coli* populations coexposed to Strep and pesticides

To couple phenotypic resistance trajectories with genetic adaptation, we carried out whole-population sequencing to identify the mutational dynamics under different selective pressures with/without pesticides. Since more populations evolved under (1/5Strep,100P) have high-level resistance compared to the other two pesticide exposure conditions, we selected the four independent populations from (1/5Strep,100P) that acquired ≥15× increase in MIC (i.e., populations 3, 5, 6, and 7) (Fig. 2D) for the whole-population sequencing every 100 generations (i.e., G0, G100, G200, G300, G400, and G500). Three parallel populations from (1/5Strep,0) (i.e., populations 1, 4, and 7) (Fig. 2A) were also sequenced as a comparison. Valid mutant alleles (one mutant allele represents one specific mutation in a gene) in the evolved populations were called out (Table S5) using standardized variant calling procedures^28^ and based on two criteria: (i) mutations leading to amino-acid-sequence change, including non-synonymous single nucleotide polymorphisms (SNPs), insertions, and deletions (INDELs), and (ii) mutant allele frequency in the population was larger than 10% at least at one time point. To have a comprehensive profile of mutational dynamics, we retrieved the frequencies of valid mutant alleles along the evolutionary path from G100 to G500.

We observed distinctive differences in mutational dynamics between populations with and without pesticide co-stressors, which corresponded to the different phenotypic Strep resistance trajectories. At an early stage in the evolved populations coexposed to (1/5Strep,100P), mutant alleles of certain genes emerged at G100 or G200, including *nuoG* (stop-gained mutations: Glu10* and Ser548*, * represents a stop codon), *nuoL* (Leu297fs, “fs” represents frameshifting), *glnE* (Ala423Val), *yaiW* (various SNPs), and *sbmA* (various SNPs), whose abundances continued to increase and became dominant after G200 or G300 (Fig. 3B). In line with that, there was a slight increase in population MICs (2 – 6×) (Fig. 3B). The populations exposed to (1/5Strep,0), which also developed mild resistance (1.5 – 5× increase in population MICs), acquired mutant alleles in the same or similar genes, such as *yaiW*, and *sbmA* (Fig. 3A). For example, the same *yaiW* mutant alleles as in (1/5Strep,100P)-7 were detected in all three sequenced (1/5Strep,0) populations.

**Fig. 3.**
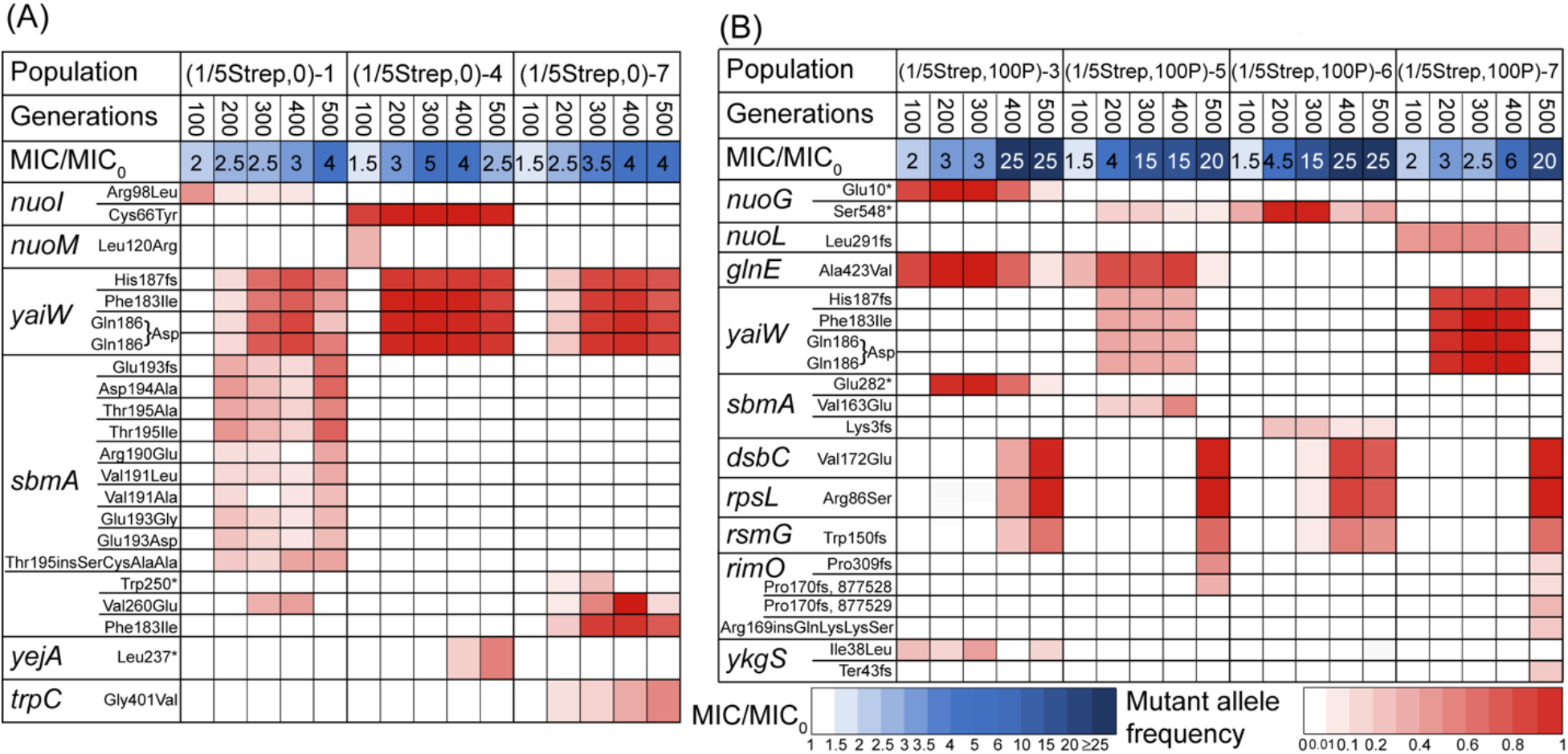
All mutated genes, the frequency of mutant alleles in the sequenced populations from G100 to G500, and the corresponding MICs. [A: 3 parallel populations exposed to (1/5Strep,0); B: 4 parallel populations exposed to (1/5Strep,100P)].

The above mutations occurred in genes encoding proteins that are not targeted by Strep, known as off-target mutations. Those genes are involved in: (i) electron transport (i.e., *nuoG* and *nuoL*), (ii) membrane permeability and transport (i.e., *yaiW* and *sbmA*), (iii) metabolism (i.e., *glnE*) and (iv) phage (i.e., *ykgS*). We then isolated resistant clones carrying these mutations. The off-target mutations in *nuoG, glnE, sbmA*, and *yaiW* all resulted in a mild (3 – 4×) increase in Strep resistance (Fig. 4, Fig. S2). Strep belongs to aminoglycoside antibiotics, which are cationic molecules. Its uptake depends on electron transport through quinones and a high membrane potential^36^. The *nuoG* mutations have been reported to reduce the uptake of Strep^10^, leading to Strep resistance. The *nuoG* mutations identified in our study were stop-gaining mutations, which may lead to loss of function. In addition, *yaiW* and *sbmA*, two closely located genes in the *E. coli* genome, encode outer membrane proteins related to antimicrobial peptide transportation^37^. Mutations in *sbmA* have been detected previously in resistant *E. coli* mutants developed under gradually increasing selective pressures of aminoglycosides^38^. The mutations in *sbmA* may result in reduced membrane permeability^39^, thus reducing the uptake of Strep. The *yaiW* mutations have not been identified in previous studies. Given the similar gene function of *yaiW* and *sbmA, yaiW* mutations likely had the same resistance mechanism as *sbmA* mutations. So far, there have not been any reports on links between Strep resistance and *glnE* mutations. The *glnE* mutation could be a resistance mutation but also could be a compensatory mutation, which could reduce the fitness cost of other mutations in the mutants.

**Fig. 4.**
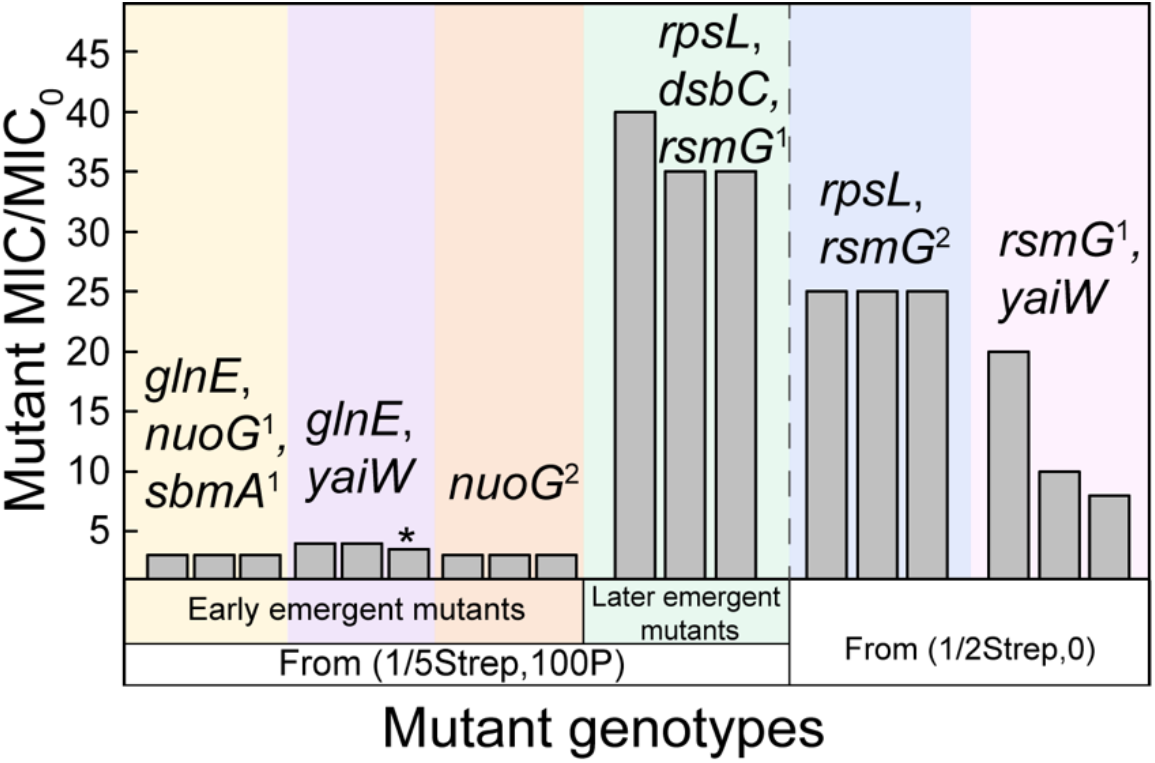
MICs of Strep-resistant mutants with different genotypes. For each genotype, three mutants were investigated; mutants containing *glnE* (Ala423Val), *nuoG*^1^ (Glu10*), and *sbmA*^1^ (Glu282*) mutations were isolated from population (1/5Strep,100P)-3 at G300; mutants containing the same *glnE* mutation and *yaiW* mutations (Phe183Ile, Gln186Asp, His187fs) were isolated from (1/5Strep,100P)-5 at G400 [Note: the mutant with “*” also had the *aidB* mutation (Pro159Gln)]; mutants containing *nuoG*^2^ (Ser548*) mutation were isolated from (1/5Strep,100P)-6 at G300; mutants with *rpsL* (Arg86Ser), *dsbC* (Val172Glu), and *rsmG*^1^ (Trp150fs) mutations were isolated from (1/5Strep,100P)-3 at G500; and mutants with the same *rpsL* and *rsmG*^2^ (Ser15insGly) mutations were isolated from (1/2Strep,0)-1 at G200).

As some of those mutations were all detected with high frequencies (80 – 100%), they seemed to accumulate in the same mutants in the evolved population. The accumulation of mutations was observed in both (1/5Strep,0) populations and early-stage (1/5Strep,100P) populations, for example, *nuoI, yaiW*, and *sbmA* in populations (1/5Strep,0)-4 and 7, as well as *nuoG, glnE*, and *sbmA* in (1/5Strep,100P)-3 at G200 (Fig. 3). The combination of mutations failed to significantly increase in MIC of those populations. For example, the population dominated by a combination of *nuoG, sbmA*, and *glnE* mutations only showed 3× MIC_0_. This result was corroborated by the MICs of resistant clones carrying *nuoG, sbmA*, and *glnE* mutations, which were also 3× MIC_0_ (Fig. 4, Fig. S2).

Although populations exposed to (1/5Strep,0) and (1/5Strep,100P) shared the same or similar trajectories of phenotypic and genotypic resistance before G300, the (1/5Strep,100P) populations exhibited a distinct succession of mutant alleles afterward. The mutations in genes *rpsL* (Arg86Ser), *rsmG* (Trp150fs), and *dsbC* (Val172Glu) occurred in all replicated (1/5Strep,100P) populations after G300. As the new mutations appeared, Strep resistance of those populations increased substantially (15 – 25×) (Fig. 3B). In the meanwhile, the early-emergent mutant alleles started to fade or completely disappeared in the populations (Fig. 3B). In contrast, the (1/5Strep,0) populations did not develop any mutations in *rpsL, rsmG*, or *dsbC* during the studied evolution period (Fig. 3A).

The *rpsL* gene encodes a Strep target protein (30S ribosomal protein S12). Mutations in *rpsL* have been frequently reported in resistant clones under strong Strep selection (e.g., a lethal dose of Strep)^10,11,40^. For example, the same *rpsL* mutation (Arg86Ser) was reported in an *E. coli* K-12 strain exposed to increasing Strep concentrations (close to and above MIC_0_) along the evolutionary path^11^. *rpsL* mutations may cause structure alteration of the target protein and reduction of Strep binding affinity. In our study, the same genetic mutation in *rpsL* was also induced under 1/2MIC_0_ Strep selection (Table S5), implying that the selection strength of (1/5Strep,100P) is as strong as 1/2MIC_0_ Strep. The resistant clones carrying that *rpsL* mutation, which were isolated from populations under both selection conditions, conferred strong phenotypic resistance (25 – 40×) (Fig. 4). More importantly, the same *rpsL* mutation was also developed under the coexposure to (1/5Strep,10P) (Table S6), suggesting pesticides, as co-stressors, could impose a strong selective pressure even at a concentration ∼10× lower than 1/5MIC of Strep.

Similar to *rpsL, rsmG* (*gidB*) encodes a methyltransferase involved in the methylation of the 16S rRNA. The loss of RsmG activity could dwindle the binding of Strep to the 30S subunit, leading to Strep resistance^41,42^. The loss-of-function mutations in *rsmG* have been identified in the previous studies^10,41^, which conferred mild to strong Strep resistance. The *rsmG* mutation in (1/5Strep,100P) populations was a 25bp frameshift deletion, which likely compromised the enzyme activity, thus resulting in Strep resistance. The same deletion mutation was also detected in mutants from the (1/2Strep,0) population (Table S5), which showed an 8 – 20× increase in MIC (Fig. 4). The other identified mutations in that population were only in *yaiW*. As the same *yaiW* mutations in the (1/5Strep,0) populations only resulted in mild resistance, the strong Strep resistance (i.e., 20× MIC_0_) in the (1/2Strep,0) population (Table S7) was more likely caused by the 25bp frameshift deletion in *rsmG*. Another mutation in *rsmG* (Ser15insGly) was developed together with the same *rpsL* mutation in another (1/2Strep, 0) population (Table S5). It might also compromise the function of RsmG, leading to Strep resistance.

The *dsbC* mutation co-occurred with the *rpsL* mutation at the same frequencies in the (1/5Strep,100P) populations after G300 (Fig. 3B), as well as in the other coexposed populations that acquired strong resistance (Table S6), suggesting that the two mutations were developed in the same genomes. Furthermore, among the randomly picked ten isolates from coexposed populations, all isolates that carried the *rpsL* mutation also contained the *dsbC* mutation (Table S8), demonstrating that *rpsL* and *dsbC* were coevolved. More interestingly, the co-selection of *rpsL* and *dsbC* mutations was not observed in populations exposed to 1/2MIC_0_ Strep (Table S5). It suggests that the coexposure of pesticide with 1/5MIC_0_ Strep can induce the emergence of specific genetic mutations, which may not show up under Strep-only selection. Mutations in *dsbC* were not linked to antibiotic resistance previously. The function of this mutation could either contribute to the resistance or reduce the fitness cost of costly resistance mutations. The isolated resistant clones with *rpsL, dsbC*, and *rsmG* mutations in (1/5Strep,100P) populations exhibited a significantly (*p* < 0.01) stronger resistance than that of resistant clones isolated from (1/2Strep,0), which also carried mutations in *rpsL* and *rsmG* (Fig. 4). Given the similar levels of resistance conferred by the two loss-of-function mutations of *rsmG*, the *dsbC* mutation was likely involved in the resistance elevation.

A significant increase in the phenotypic resistance showed up right after the first emergence of the *rpsL* + *dsbC* + *rsmG* mutant in the (1/5Strep,100P) populations. We used a titration experiment to determine the lowest fraction of strongly resistant mutation that could cause a change in the phenotypic resistance of the population. We constructed mock populations containing the strongly resistant *rpsL* + *dsbC* + *rsmG* mutant and the wild type at different ratios. We observed that even when the fraction of the strongly resistant mutant was lowered to 10^−3^, the population MIC still increased significantly to 15× MIC_0_ (Fig. 5). A fraction as low as 10^−5^ still caused a five-fold increase in MIC of the population. This result could explain the high-level resistance obtained in the (1/5Strep,100P)-5 population at G300 and G400 (Fig. 3B), where mutations conferring strong resistance were not identified. The strongly resistant mutants might have already arisen in the populations, which led to the substantial increase in the population MIC, but the mutation frequency could be too low (i.e., < 1/sequencing depth ∼ 1/800) to be detected by the whole-population sequencing.

**Fig. 5.**
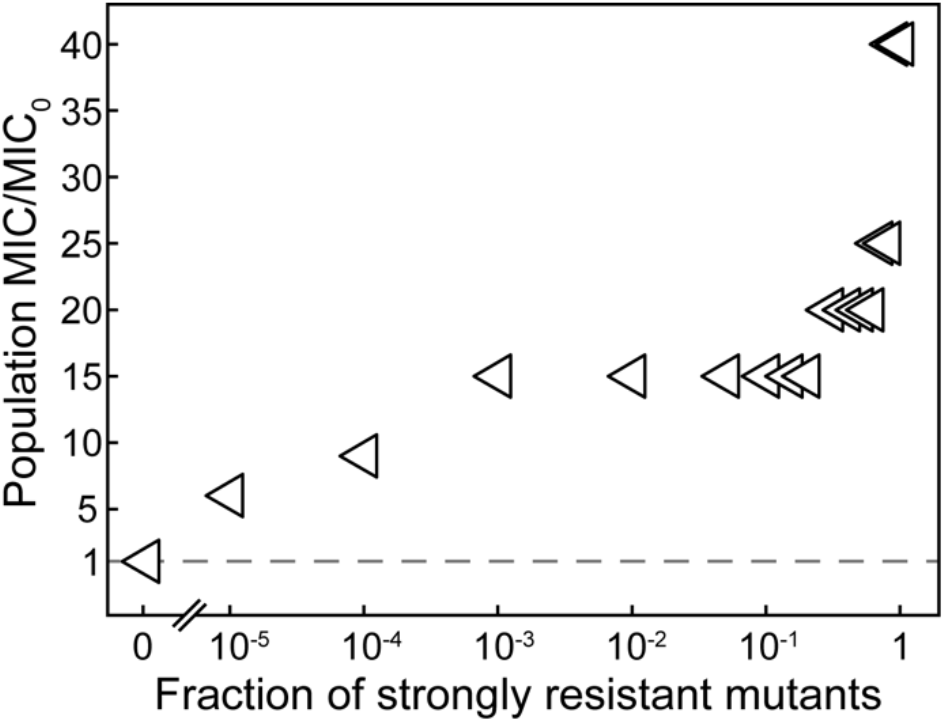
MICs of mock populations containing the wild type and various fractions of the strongly resistant mutant carrying the *rpsL, dsbC*, and *rsmG* mutations.

### Fitness evolutionary trajectories in *E. coli* populations under the coexposure

The relative fitness of resistant mutants in a population determines their evolutionary directions with or without the selective pressure. For example, if the resistant mutants had higher fitness than the wild type in the presence of the selective pressure, the wild type would be outcompeted and toward extinction in the population, leading to the persistence of phenotypic resistance. According to the succession pattern in the coexposed populations, the late emergent mutants with strong Strep resistance seem to have the highest fitness, followed by the early emergent mutants, whereas the wild type has the lowest fitness under the selective pressure of (1/5Strep,100P). This was corroborated by competition tests between the wild type, the mildly resistant mutant (with *nuoG, glnE*, and *sbmA* mutations), and the strongly resistant mutant (with *rpsL, dsbC*, and *rsmG* mutations). Expectedly, starting at 1% in the population, both the mildly and strongly resistant mutants outcompeted the wild type cells under the selective pressure (1/5Strep,100P) after 50 generations (Fig. 6), indicating higher growth fitness of those resistant mutants than the wild type. Additionally, the strongly resistant mutant outcompeted the mildly resistant mutant in constructed cocultures coexposed to (1/5Strep,100P), consistent with the succession observed in the actual coexposed populations. The same result (Fig. S3) was observed when the strongly resistant mutant was competing with the other two early emergent mutants, which were isolated from the coexposed populations and conferred mild resistance. The results of competition tests under different selective pressures reveal that the presence of Strep is the major factor maintaining the growth advantage of preexisting resistant mutants, whereas pesticides did not play a role (Table S9). We further estimated the minimal selective concentrations (MSC) of Strep for these resistant mutants by competing them with wild-type cells. There was variation in the MSC of different mutants; the lowest MSC corresponded to 1/40 MIC_0_ (i.e., 200 μg/L) for the late emergent mutant with strong resistance, while the highest MSC was 1/10 MIC_0_ for the mutant with a combination of *nuoG, glnE*, and *sbmA* mutations (Table S10). These results led to the prediction that enrichment of the preexisting resistant mutants is possible at Strep concentrations significantly below the MICs of wild-type strains, and with a lower MSC the strongly resistant mutant could even be more favorably selected over the mildly resistant mutants in a population.

**Fig. 6.**
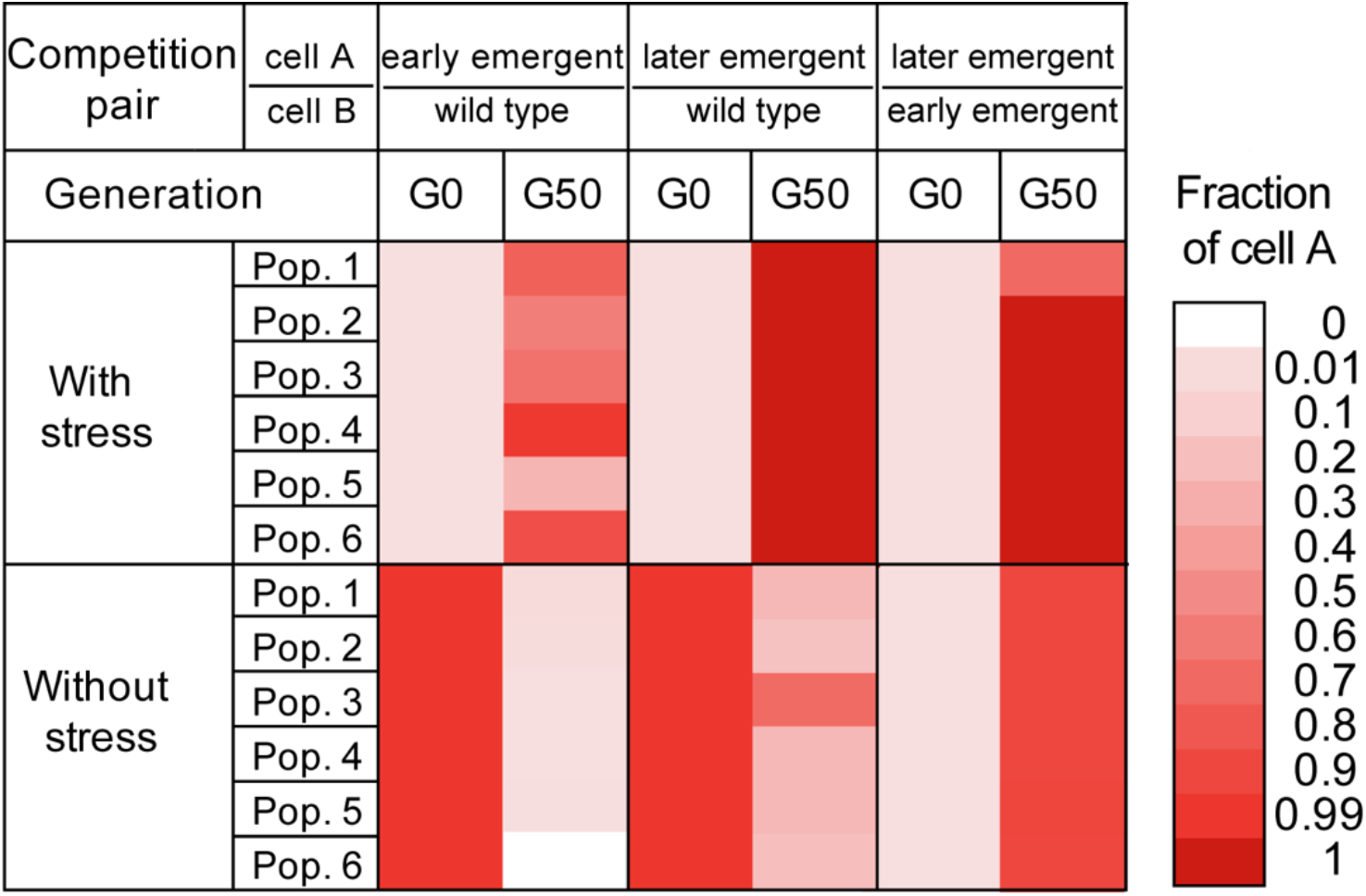
Growth competition between the wild type, the early emergent mutant with mild Strep resistance (carrying *nuoG, glnE, and sbmA* mutations), and the late emergent mutant with strong Strep resistance (carrying *rpsL, dsbC*, and *rsmG* mutations) in LB medium with and without the selective pressure (1/5Strep,100P) (six parallel populations containing cell A and B were performed; two initial fractions of cell A, i.e., 1% and 99%, were included).

Resistant mutants usually have fitness costs and are likely to become extinct in competition with wild-type cells in the absence of selective pressures, leading to the reversal of antibiotic-resistance evolution. To test the reversibility of the evolution direction after the removal of the selective pressure, we conducted competition tests starting with 99% of the mildly or strongly resistant mutants and 1% of wild-type cells in LB broth without any selective pressure. As expected, when competing against the wild type without the selective pressure, both mildly and strongly resistant mutants lost their growth advantages and were outcompeted after 50 generations (Fig. 6, Fig. S3). Interestingly, in competitions between mildly and strongly resistant mutants without the selective pressure, the strongly resistant mutant still outcompeted the mildly resistant mutant even when the relative abundance of the strongly resistant mutant was as low as 1% (Fig. 6, Fig. S3). It demonstrated a higher fitness of the strongly resistant mutant than the mildly resistant mutant regardless of the selective pressure, indicating that the evolution direction may not be reversed by simply removing the selective pressure in a population where the two mutant types coexist. We further demonstrated this by growing (1/5Strep,100P)-5 (G500) population in LB medium without the selective pressure. After 50 generations, the fraction of strongly resistant mutant with *rpsL* + *dsbC* + *rsmG* mutations remained at ∼100% in the population (Table S11). We also explored the reversibility of resistance evolution in populations from the selection of 1/2MIC_0_ Strep. The fraction of the strongly resistant mutant dropped from nearly 100% to 5% in (1/2Strep,0)-1 after 50 generations (Table S11). These results indicate that the genetic backgrounds of the populations that evolved under both exposure conditions are quite different, which affected the reversibility of the evolution direction.

We then compared the relative fitness of resistant mutants from (1/5Strep,100P) populations with those from Strep only conditions. The more positive the selection coefficient (A/B) is the higher fitness cell A has over cell B, and vice versa. From the results, we could draw two conclusions. First, the mutant with the highest fitness came from the (1/2Strep,0) condition at G200 with *rsmG* + *yaiW* mutations, followed by the mutant from the (1/5Strep,100P) condition at G500 with *rpsL* + *dsbC* + *rsmG* mutations. The mutant with the most significant fitness defect was the resistant clone from (1/5Strep,0)-1 at G500 (Fig. 7). Usually, it has been known that the stronger the selection strength, the lower the fitness of the obtained resistant mutants^12,43^. However, our results strongly fail to support this perspective. On the contrary, we observed mutants with a higher fitness from the stronger selection conditions, whereas the most defective mutant evolved from the weakest selection condition. This implies that the fitness of resistant mutants might be dependent on their characteristics with regard to specific mutations, not the strength of selection. Second, the relative fitness of *rpsL* + *dsbC* + *rsmG* mutant is slightly higher than *rpsL* + *rsmG* mutant, suggesting the role of *dsbC* mutation in reducing the fitness cost caused by resistance mutations (e.g., *rpsL* mutation).

**Fig. 7.**
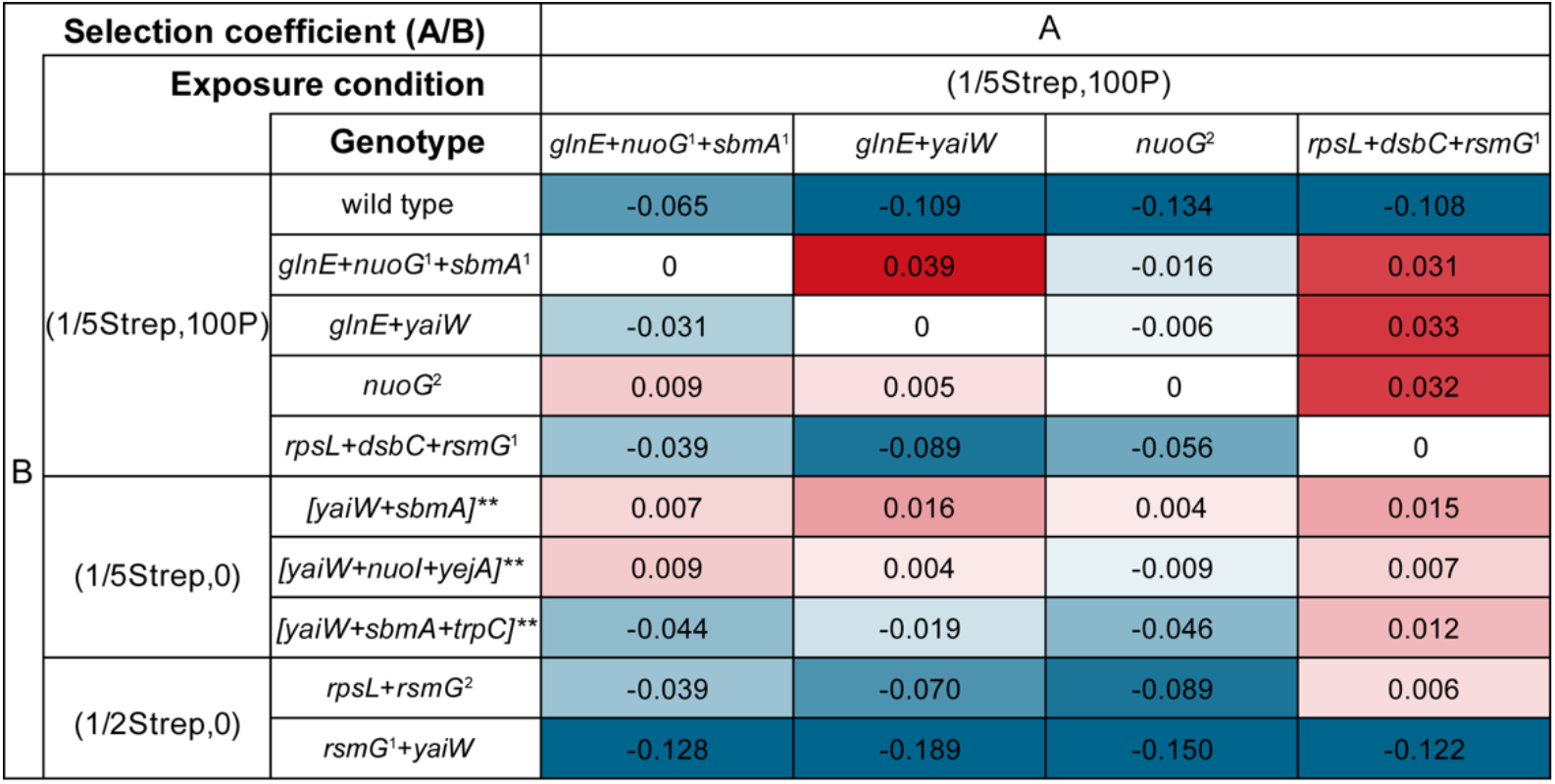
Selection coefficients of cells with different genotypes isolated from (1/5Strep,100P) populations and mutants from Strep only selection conditions. (mutations: *glnE* (Ala423Val), *nuoG*^1^ (Glu10*), *sbmA*^1^ (Glu282*), *yaiW* (Phe183Ile, Gln186Asp, His187fs), *nuoG*^2^ (Ser548*), *rpsL* (Arg86Ser), *dsbC* (Val172Glu), *rsmG*^1^ (Trp150fs), and *rsmG*^2^ (Ser15insGly); []** indicates the suspect genotypes of the mutant based on the whole-population sequencing results).

## Discussion

Selective pressures are of great importance to understand the evolution of antibiotic resistance in the environment. Previous studies have demonstrated that sub-MIC antibiotics can serve as selective pressures and facilitate the evolution of antibiotic resistance. However, very little is understood about how non-antibiotic co-stressors could affect the antibiotic resistance selection and the phenotypic, genomic, and fitness evolutionary trajectories. Here, we focused on the evolution of antibiotic resistance in microbial populations under the coexposure of sub-MIC (i.e., 1/5MIC_0_) Strep and the co-stressor, pesticides. We first showed that the exposure to pesticides, in addition to sub-MIC Strep, led to the selection of strong resistance, which could not be driven by the Strep-only exposure. Next, we coupled the evolutionary trajectories of the phenotypic resistance with the evolutionary trajectories of genotypes and growth fitness. We observed the succession of dominant mutants in the coexposed populations from the off-target mutations to the target-modification mutations during the evolution. This displacement pattern led to the transition from mild to strong phenotypic resistance. Compared to the mildly resistant mutants, the strongly resistant mutants developed under the coexposure exhibited higher growth fitness with and without the selective pressure, as well as a lower MSC, which could favor the proliferation and sustain the dominance of strongly resistant mutants under environmentally relevant conditions.

There are several implications of these results. First, the role of pesticide co-stressors in promoting the selection of strong antibiotic resistance would be of greater concern, which has been largely overlooked. The effective concentration of pesticides to exhibit the synergistic effect was as low as 20 µg/L in total, which is typical in agricultural and wastewater-related environments, where antibiotics could also exist at low levels^21,44,45^. Thus, such environmental conditions may select for *de novo* mutants with much stronger resistance than those selected by the same level of antibiotics alone. Pharmaceutical co-stressors examined in this study did not perform in the same way as pesticides, suggesting the specificity of pesticides as co-stressors to promote antibiotic resistance selection. More co-stressors are worth further investigation in future studies, as well as the effect of individual pesticides. In addition, we demonstrated that pesticides are not involved in the selection of preexisting mutants. The selection of preexisting mutants conferring resistance is solely dependent on the antibiotic, and minimal selective concentrations of the obtained mutants could be as low as 1/40 MIC_0_. This result is consistent with the previous study, which suggests the selection of preexisting mutants could be much lower than inhibitory concentrations^7^.

Second, our results showed various mutations leading to Strep resistance, including some novel resistance mechanisms. Target-modification mutations like *rpsL* mutations and *rsmG* mutations have been reported to cause strong resistance to Strep^10,11,43^. However, off-target mutations conferring resistance are less known. In this study, by studying the evolutionary trajectories of genomic evolution, we have identified novel off-target mutations conferring mild Strep resistance. For example, the mutations in *yaiW* and *sbmA* genes, which encode peptide antibiotic transporters on cell membranes, might reduce membrane permeability^39^, and thus lower the uptake of Strep. Notably, the accumulation of off-target mutations failed to lead to strong resistance. This result disagrees with the previous study, where the combination of five mutations led to a significantly high level of resistance, although individual mutations did not confer strong resistance^10^. It might be due to the difference in acquired mutation spectra, exposure length (900 vs. 500 generations), and tested bacterial species (*S. enterica* vs. *E. coli*) between the two studies. Moreover, our results implied that the *dsbC* mutation, which was coevolved with *rpsL* under the coexposure, could elevate Strep resistance and lower the fitness cost caused by other resistance mutations. These results need to be followed with mechanistic studies to determine the precise functional roles of the mutations. Next, the prevalence of these mutations evolved from laboratory systems needs to be examined in the environments potentially impacted by antibiotics and pesticides. If the resistance mutations could be identified in real environments, they could serve as biomarkers indicative of antibiotic resistance.

Third, effective mitigation strategies against the development and propagation of strong antibiotic resistance should be more carefully and comprehensively made considering selective pressures, genetic background, and subgroup fitness comparison in microbial populations under varying environmental conditions. The evolutionary trajectories in this study highlighted the succession from genotypes conferring mild resistance to those conferring strong resistance in coexposed populations, which was not observed in populations exposed to sub-MIC Strep only. Removal of stressors is one strategy to control the development and/or proliferation of strongly resistant mutants, hence reducing antibiotic resistance in a population. However, it might not work effectively for the populations that have developed strong resistance from a transitional phase dominated by mildly resistant mutants with lower fitness. Moreover, since some resistant mutants likely have an MSC much lower than MIC, high removal efficiencies of antibiotics would be needed to avoid the selection of antibiotic-resistant mutants. Collectively, the presence of antibiotics and other co-stressors in specific environments makes it more challenging to control antibiotic resistance. In this case, pesticides and antibiotics need to be effectively removed at an early stage of exposure or even before they enter the receiving environment, for example, enhancing their removal efficiencies in wastewater treatment plants.

### Data Availability

All the whole-population/genome sequencing data have been deposited in the NCBI SRA database under Accession No. PRJNA605244.

## Acknowledgements

We would like to give thanks to Chris L. Wright at the Roy J. Carver Biotechnology Center, University of Illinois at Urbana–Champaign for whole-population sequencing support.

