## Supplementary Information for "Specific phenotypic, genomic, and fitness evolutionary trajectories toward antibiotic resistance induced by pesticide co-stressors in *Escherichia coli*"

### Supplementary Methods

#### Measuring growth rate under different conditions

Growth rates were measured at 37 °C in non-selective LB broth using Synergy 2 Multi-Mode Microplate Reader (BioTek Instruments). Each population exposed to (0,0), (1/5Strep,0), (1/5Strep,100P), and (0,100P) had 10 biological replicates on a 96-well plate. The inoculum was the culture grown overnight from 10 μL of 2× diluted archived population in 2 mL LB broth. The 96-well plate was incubated in the Microplate Reader for 24 h with continuous shaking at 150 rpm in the dark, and the optical density at 600 nm ($OD_{600}$) was measured every 15 min. The max growth rates and lag time were calculated by the software Gen 5.

#### SNP genotyping assays

We designed PCR-based SNP genotyping assays via Custom TaqMan SNP Genotyping Assays (Thermo Fisher Scientific) targeting six parallel mutant alleles: *nuoG*$^1$ (Glu10*), *nuoG*$^2$ (Ser548*), *glnE* (Ala423Val), *sbmA*$^1$ (Glu282*), *rpsL* (Arg86Ser), and *dsbC* (Val172Glu). The genotyping reactions were carried out in a total volume of 5 μL (2.75 μL master mix + customized genotyping assay and 2.25 μL diluted DNA sample of 3 ng/μL) according to the manufacturer's instructions. The assays were performed in 96-well plates on a real-time PCR instrument QuantStudio 3 (Thermo Fisher Scientific), following the recommended thermal cycling conditions. Thermo Fisher Cloud "Genotyping" application was used to generate allele calls.

The SNP genotyping assay was used to confirm genotypes of isolated mutants and also adapted to estimate (i.e., semi-quantify) the mutant allele frequencies in non-sequenced coexposed populations along the evolutionary path and in constructed cocultures for the competition tests. The quantification standards were prepared by mixing gDNA of cell A with gDNA of cell B at different ratios, resulting in the fraction of the mutant alleles at 1%, 5%, 10%, 15%, 20%, 30%, 40%, 50%, 60%, 70%, 80%, 90%, 95%, and 99%. The fraction of mutant alleles in the tested populations was estimated by comparing the allelic discrimination plot to those of the standard mixtures (Fig. S1).

**Determination of minimal selective concentrations (MSC)**

The MSC was estimated as the streptomycin concentration where the resistant mutant has an equal growth rate with the wild type. We set up competition tests between a resistant mutant and the wild type (1:1) with Strep addition ranging from 0 to 1/2 $MIC_0$ (0, 1/100, 1/80, 1/40, 1/20, 1/10, 1/8, 1/5, 1/2). The MSC was determined as the lowest concentration of Strep, at which the fraction of the resistant mutant did not decrease after the competition test.

**Table S1** Environmental concentrations of selected pesticides and pharmaceuticals

| Name | Classification | Conc. (μg/L) | Environmental samples and references | Purchase |
|---|---|---|---|---|
| | | **Pesticides** | | |
| 2,4-D | Herbicide | 0.2 | Urban run-off[1] | Sigma |
| Atrazine | Herbicide | 0.5 | Groundwater and surface water[2] | Sigma |
| Benomyl | Fungicide | 0.2 | Surface water[3] | Sigma |
| Carbaryl | Insecticide | 4.8 | Surface water[4] | Sigma |
| Carbofuran | Insecticide | 0.38 | Ground and surface water[5] | Sigma |
| Chlorpyrifos | Pesticide | 0.4 | Lake[6] | AK Scientific |
| Clotrimazole | Fungicide | 0.1 | Wastewater[7] | AK Scientific |
| DEET | Biocide | 3 | Wastewater influent[8] | Sigma |
| Diazinon | Insecticide | 0.3 | Wastewater[9] | Sigma |
| Diuron | Herbicide | 1 | Urban run-off[1] | AK Scientific |
| Fipronil | Insecticide | 0.2 | Urban surface water[10] | AK Scientific |
| Imazalil | Fungicide | 0.4 | River[11] | AK Scientific |
| Imidacloprid | Insecticide | 0.4 | Ground and surface water[5] | Sigma |
| Irgarol | Biocide | 0.2 | Coastal water[12] | Sigma |
| Linuron | Herbicide | 2 | Rivers[13] | Sigma |
| Mecoprop | Herbicide | 2 | Urban run-off[1] | Sigma |
| Metaldehyde | Pesticide | 0.5 | Surface water[14] | Sigma |
| Metolachlor | Herbicide | 0.4 | Wastewater[9] | Sigma |
| Propiconazole | Fungicide | 1 | Wastewater[15] | Sigma |
| Tebuconazole | Fungicide | 0.5 | Wastewater[16] | Sigma |
| Terbuthylazine | Herbicide | 0.65 | Ground and surface water[5] | AK Scientific |
| Terbutryn | Herbicide | 0.5 | Rivers[17] | Sigma |
| Thiabendazole | Fungicide | 0.2 | Wastewater influent[18] | Sigma |
| **Total** | | **19.83** | | |
| | | **Pharmaceuticals** | | |
| Paracetamol | | 5 | Wastewater[8] | Sigma |
| Gabapentin | | 5 | Wastewater[8] | AK Scientific |
| Ibuprofen | | 2 | Wastewater[19] | Sigma |
| Atenolol | | 4 | Wastewater[20] | Sigma |
| Caffeine | | 5 | Wastewater[8] | Sigma |
| Amantadine | | 1 | Wastewater[21] | AK Scientific |
| Fluconazole | | 0.5 | Wastewater[22] | AK Scientific |
| Flucytosine | | 1 | Wastewater[8] | AK Scientific |
| Carbamazepine | | 2 | Wastewater[8] | Sigma |
| Ranitidine | | 2 | Wastewater[8] | Sigma |
| O-Desmethylvenlafaxine | | 1 | Wastewater[8] | Sigma |
| **Total** | | **28.5** | | |

**Table S2** Output of max V and lag time under different selection pressure

| (0,0) | | (1/5Strep,0) | | (1/5Strep,100P) | | (0,100P) | |
|---|---|---|---|---|---|---|---|
| **Max V** | **Lag time** | **Max V** | **Lag time** | **Max V** | **Lag time** | **Max V** | **Lag time** |
| 5.864 | 2:15:15 | 5.446 | 2:13:16 | 5.708 | 2:17:42 | 5.651 | 2:07:23 |
| 5.179 | 2:06:50 | 5.759 | 2:11:40 | 5.477 | 2:10:30 | 5.486 | 2:06:06 |
| 6.073 | 2:04:07 | 5.099 | 2:04:36 | 5.727 | 2:08:20 | 5.632 | 1:59:17 |
| 5.265 | 1:59:36 | 5.599 | 2:05:09 | 5.568 | 2:11:37 | 5.055 | 1:58:20 |
| 5.653 | 2:00:22 | 5.877 | 2:09:30 | 5.116 | 1:58:05 | 5.479 | 1:56:45 |
| 5.601 | 2:02:51 | 5.372 | 1:59:08 | 4.739 | 1:58:54 | 5.35 | 2:03:17 |
| 5.89 | 2:03:46 | 5.341 | 2:03:41 | 5.08 | 1:59:16 | 5.362 | 2:01:08 |
| 5.582 | 1:58:52 | 5.705 | 2:05:17 | 5.645 | 2:01:44 | 5.356 | 1:56:52 |
| 6.115 | 2:09:45 | 5.868 | 2:07:11 | 5.692 | 2:02:39 | 5.758 | 2:01:08 |
| 5.362 | 1:56:16 | 5.531 | 2:11:02 | 5.255 | 2:02:33 | 5.763 | 1:57:14 |
| Average | | | | | | | |
| 5.658 | 2:03:46 | 5.56 | 2:07:03 | 5.4 | 2:05:08 | 5.489 | 2:00:45 |

**Table S3** Strep resistance developed in *E. coli* populations exposed to pesticides only

| Population | MIC ($MIC_0$ = 8 mg/L) | | | |
|---|---|---|---|---|
| | (0,0) | (0,1P) | (0,10P) | (0,100P) |
| Rep-1 | 1× $MIC_0$ | 1× $MIC_0$ | 1× $MIC_0$ | 1× $MIC_0$ |
| Rep-2 | 1× $MIC_0$ | 1× $MIC_0$ | 1× $MIC_0$ | 1× $MIC_0$ |
| Rep-3 | 1× $MIC_0$ | 1.5× $MIC_0$ | 1× $MIC_0$ | 1.5× $MIC_0$ |
| Rep-4 | 1× $MIC_0$ | 1.5× $MIC_0$ | 1.5× $MIC_0$ | 1.5× $MIC_0$ |
| Rep-5 | 1× $MIC_0$ | 1.5× $MIC_0$ | 1.5× $MIC_0$ | 1.5× $MIC_0$ |
| Rep-6 | 1× $MIC_0$ | 1.5× $MIC_0$ | 1.5× $MIC_0$ | 1.5× $MIC_0$ |
| Rep-7 | 1× $MIC_0$ | 1.5× $MIC_0$ | 1.5× $MIC_0$ | 1.5× $MIC_0$ |
| Rep-8 | 1.5× $MIC_0$ | 1.5× $MIC_0$ | 1.5× $MIC_0$ | 1.5× $MIC_0$ |

**Table S4** Strep resistance developed in *E. coli* populations exposed to Strep at 1/5$MIC_0$ and pharmaceuticals (Ph)

| Population | MIC ($MIC_0$ = 8 mg/L) | | | |
|---|---|---|---|---|
| | (1/5Strep,0) | (1/5Strep,1Ph) | (1/5Strep,10Ph) | (1/5Strep,100Ph) |
| Rep-1 | 4× $MIC_0$ | 4× $MIC_0$ | 4× $MIC_0$ | 4× $MIC_0$ |
| Rep-2 | 4× $MIC_0$ | 4× $MIC_0$ | 4× $MIC_0$ | 4× $MIC_0$ |
| Rep-3 | 4× $MIC_0$ | 6× $MIC_0$ | 4× $MIC_0$ | 4× $MIC_0$ |
| Rep-4 | 4× $MIC_0$ | 4× $MIC_0$ | 4× $MIC_0$ | 4× $MIC_0$ |
| Rep-5 | 6× $MIC_0$ | 4× $MIC_0$ | 4× $MIC_0$ | 6× $MIC_0$ |
| Rep-6 | 4× $MIC_0$ | 4× $MIC_0$ | 6× $MIC_0$ | 6× $MIC_0$ |
| Rep-7 | 4× $MIC_0$ | 4× $MIC_0$ | 4× $MIC_0$ | 4× $MIC_0$ |
| Rep-8 | 6× $MIC_0$ | 6× $MIC_0$ | 4× $MIC_0$ | 4× $MIC_0$ |

**Table S6** The presence of *rpsL* and *dsbC* mutations in non-sequenced coexposed populations and the corresponding MICs

| **Population** | **Generation** | ***rpsL*** | ***dsbC*** | **Population MIC** |
|---|---|---|---|---|
| (1/5Strep,10P)-5 | G100 | N.D. | N.D. | 2× $MIC_0$ |
| | G200 | 20% | 20% | 25× $MIC_0$ |
| | G300 | 20% | 20% | 25× $MIC_0$ |
| | G400 | 100% | 100% | 25× $MIC_0$ |
| | G500 | 100% | 100% | 25× $MIC_0$ |
| (1/5Strep,10P)-6 | G100 | N.D. | N.D. | 2× $MIC_0$ |
| | G200 | < 1% | < 1% | 6× $MIC_0$ |
| | G300 | 1 – 5% | 1 – 5% | 9× $MIC_0$ |
| | G400 | N.D. | N.D. | 8× $MIC_0$ |
| | G500 | N.D. | N.D. | 8× $MIC_0$ |
| (1/5Strep,10P)-7 | G100 | N.D. | N.D. | 1.5× $MIC_0$ |
| | G200 | N.D. | N.D. | 3.5× $MIC_0$ |
| | G300 | N.D. | N.D. | 3.5× $MIC_0$ |
| | G400 | 1% | 1% | 15× $MIC_0$ |
| | G500 | 100% | 100% | 20× $MIC_0$ |
| (1/5Strep,100P)-4 | G100 | N.D. | N.D. | 2× $MIC_0$ |
| | G200 | N.D. | N.D. | 3× $MIC_0$ |
| | G300 | N.D. | N.D. | 3× $MIC_0$ |
| | G400 | N.D. | N.D. | 3.5× $MIC_0$ |
| | G500 | 1% | 1% | 9× $MIC_0$ |

Note: N.D.: not detected; all non-sequenced populations shown in Fig. 2 were tested and only populations that contain the *rpsL* and *dsbC* mutations are listed in this table.

**Table S7** Strep resistance developed in *E. coli* populations exposed to Strep only at 1/2 $MIC_0$ (1/2Strep,0)

| **Population** | **(1/2Strep,0) at 50G** | **(1/2Strep,0) at 100G** | **(1/2Strep,0) at 200G** |
|---|---|---|---|
| Rep-1 | ≥20× $MIC_0$ | ≥20× $MIC_0$ | ≥20× $MIC_0$ |
| Rep-2 | 3× $MIC_0$ | 5× $MIC_0$ | 15× $MIC_0$ |
| Rep-3 | 3× $MIC_0$ | 3× $MIC_0$ | ≥20× $MIC_0$ |
| Rep-4 | 2× $MIC_0$ | 3× $MIC_0$ | 3× $MIC_0$ |
| Rep-5 | 2× $MIC_0$ | 5× $MIC_0$ | 6× $MIC_0$ |
| Rep-6 | 2× $MIC_0$ | 3× $MIC_0$ | 3× $MIC_0$ |
| Rep-7 | 2× $MIC_0$ | 3× $MIC_0$ | 10× $MIC_0$ |
| Rep-8 | 3× $MIC_0$ | ≥20× $MIC_0$ | ≥20× $MIC_0$ |

**Table S8** The detection of *rpsL* and *dsbC* mutations in 10 randomly picked isolates from (1/5Strep,100P)-3 and (1/5Strep,10P)-5 populations using non-selective LB agar plates

| **Population** | **Frequency of *rpsL* and/or *dsbC* mutants** | **Genotypes in each mutant** |
|---|---|---|
| (1/5Strep,100P)-3-G100 | 0/10 | N.D. |
| (1/5Strep,100P)-3-G200 | 0/10 | N.D. |
| (1/5Strep,100P)-3-G300 | 0/10 | N.D. |
| (1/5Strep,100P)-3-G400 | 3/10 | *rpsL* & *dsbC* |
| (1/5Strep,100P)-3-G500 | 9/10 | *rpsL* & *dsbC* |
| (1/5Strep,10P)-5-G100 | 0/10 | N.D. |
| (1/5Strep,10P)-5-G200 | 2/10 | *rpsL* & *dsbC* |
| (1/5Strep,10P)-5-G300 | 2/10 | *rpsL* & *dsbC* |
| (1/5Strep,10P)-5-G400 | 10/10 | *rpsL* & *dsbC* |
| (1/5Strep,10P)-5-G500 | 10/10 | *rpsL* & *dsbC* |

N.D.: not detected

**Table S9** Competition tests under different selection conditions

| **Selection condition** | *nuoG*[1]+*glnE*+*sbmA*: wild type | | *glnE* +*yaiW*: wild type | | *nuoG*[2]: wild type | | *rpsL*+*dsbC*+*rsmG*[1]: wild type | |
|---|---|---|---|---|---|---|---|---|
| | G0 | G9 | G0 | G9 | G0 | G9 | G0 | G9 |
| No stress | 40% | 25% | 30% | 15% | 30% | 15% | 30% | 17% |
| (0,100P) | 40% | 25% | 30% | 15% | 30% | 15% | 30% | 15% |
| (1/5Strep,0) | 40% | 45% | 30% | 40% | 30% | 35% | 30% | 60% |
| (1/5Strep,100P) | 40% | 50% | 30% | 40% | 30% | 36% | 30% | 60% |

**Table S10** MSC of resistant mutants from (1/5Strep, 100P) populations

| Mutant genotype | MSC (of $MIC_0$) | MSC (absolute value μg/L) |
|---|---|---|
| *nuoG*$^1$ + *glnE* + *sbmA* | 1/10 $MIC_0$ | 800 |
| *glnE* + *yaiW* | 1/10 ~ 1/20 $MIC_0$ | 400 ~ 800 |
| *nuoG*$^2$ | 1/20 $MIC_0$ | 400 |
| *rpsL* + *dsbC* + *rsmG*$^1$ | 1/40 $MIC_0$ | 200 |

**Table S11** The change of *rpsL* mutant fraction in (1/2Strep,0)-1-G200 populations and (1/5Strep,100P)-5-G500 population after grown in LB medium without the corresponding selection pressure for 50 generations

| Population | Fraction of *rpsL* mutants before the no-stress growth | Fraction of *rpsL* mutants after the no-stress growth |
|---|---|---|
| (1/2Strep,0)-1-G200 | ~100% | 5% |
| (1/5Strep,100P)-5-G500 | ~100% | ~100% |

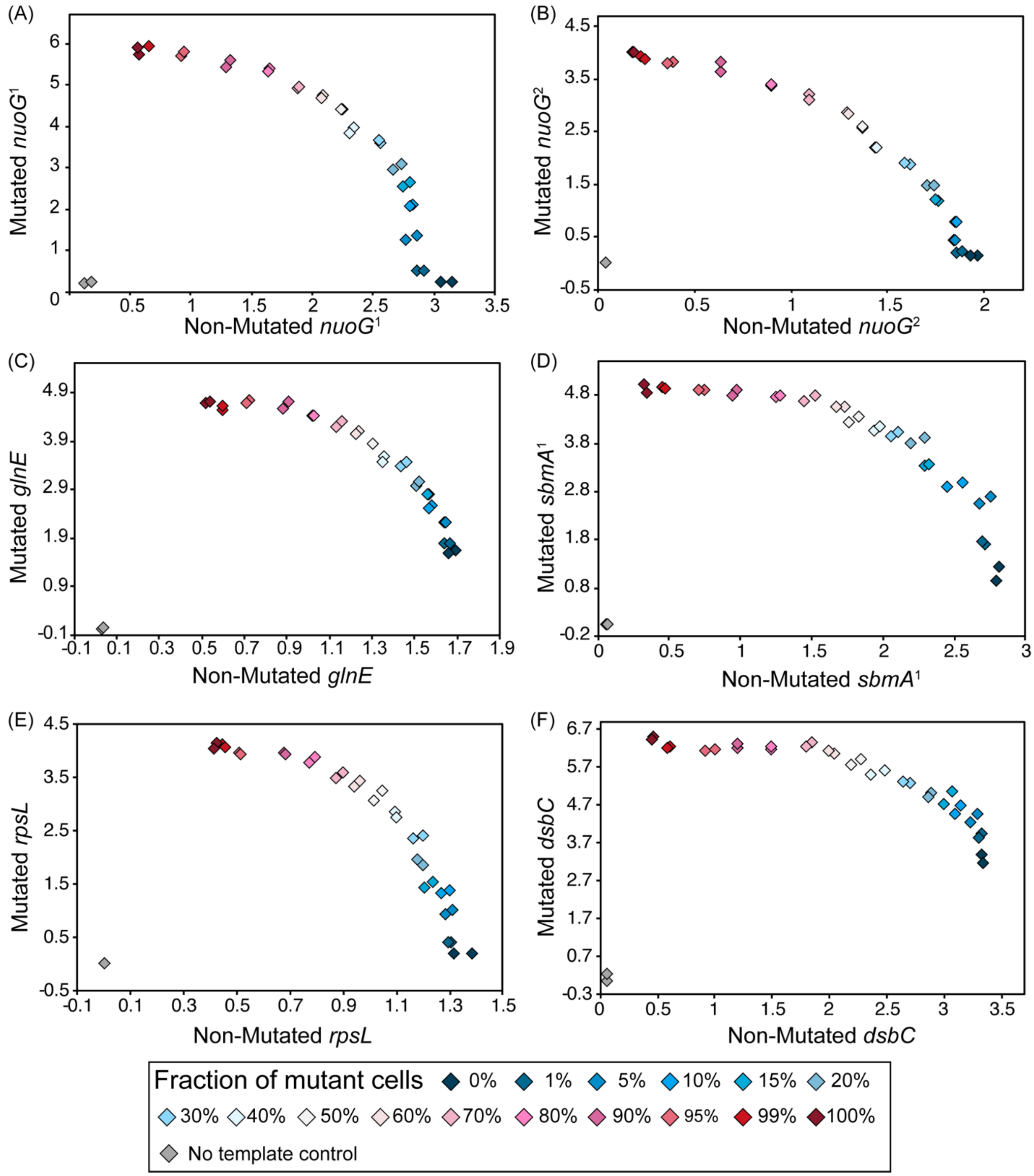


**Fig. S1.** Allelic discrimination plots obtained for *nuoG*[1] (Glu10*) (A), *nuoG*[2] (Ser548*) (B), *glnE* (Ala423Val) (C), *sbmA*[1] (Glu282*) (D), *rpsL* (Arg86Ser) (E), and *dsbC* (Val172Glu) (F) SNP mutations on standard mixtures at different fractions of mutant cells.

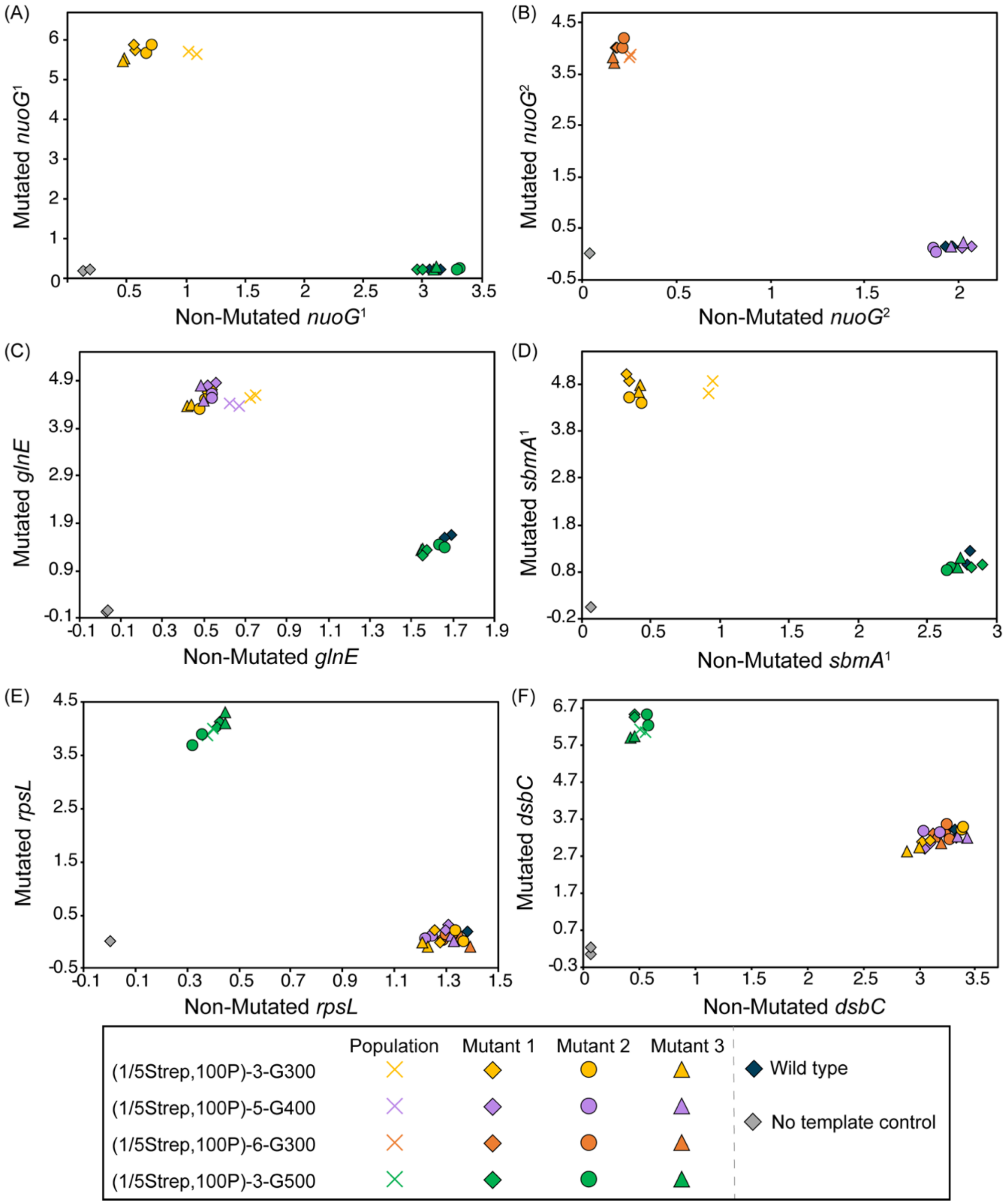


**Fig. S2.** Allelic discrimination plots obtained for *nuoG*[1] (Glu10*) (A), *nuoG*[2] (Ser548*) (B), *glnE* (Ala423Val) (C), *sbmA*[1] (Glu282*) (D), *rpsL* (Arg86Ser) (E), and *dsbC* (Val172Glu) (F) SNP mutations on isolated mutants and their originated populations.

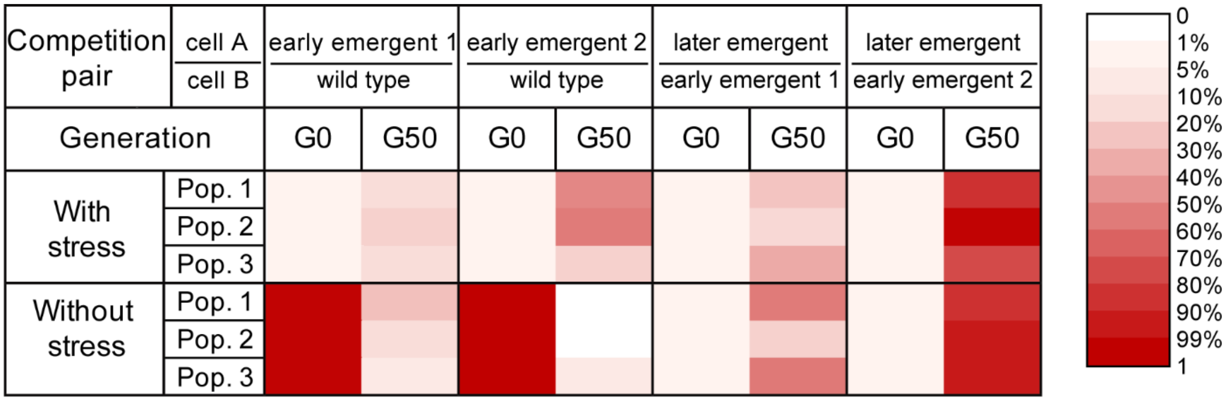


**Fig. S3.** Growth competition between the wild type, the two early emergent mutants with mild Strep resistance [early emergent 1: with *nuoG*[2] (Ser548*) mutation; early emergent 2: with *glnE* (Ala423Val) and *yaiW* (Phe183Ile, Gln186Asp, His187fs) mutations], and the late emergent mutant with strong Strep resistance, carrying *rpsL*, *dsbC*, and *rsmG*[1] (Trp150fs) mutations, in LB medium with and without the selection pressure (1/5Strep,100P) (Three parallel populations containing cell type A and B were performed with an initial fraction of cell A at 1% and 99%).
